# A-GHN: Attention-based Fusion of Multiple GraphHeat Networks for Structural to Functional Brain Mapping

**DOI:** 10.1101/2021.08.12.456134

**Authors:** Subba Reddy Oota, Archi Yadav, Arpita Dash, Raju S. Bapi, Avinash Sharma

## Abstract

Over the last decade, there has been growing interest in learning the mapping from structural connectivity (SC) to functional connectivity (FC) of the brain. The spontaneous fluctuations of the brain activity during the restingstate as captured by functional MRI (rsfMRI) contain rich non-stationary dynamics over a relatively fixed structural connectome. Among the modeling approaches, graph diffusion-based methods with single and multiple diffusion kernels approximating static or dynamic functional connectivity have shown promise in predicting the FC given the SC. However, these methods are computationally expensive, not scalable, and fail to capture the complex dynamics underlying the whole process. Recently, deep learning methods such as GraphHeat networks along with graph diffusion have been shown to handle complex relational structures while preserving global information. In this paper, we propose a novel attention-based fusion of multiple GraphHeat networks (A-GHN) for mapping SC-FC. A-GHN enables us to model multiple heat kernel diffusion over the brain graph for approximating the complex *Reaction Diffusion* phenomenon. We argue that the proposed deep learning method overcomes the scalability and computational inefficiency issues but can still learn the SC-FC mapping successfully. Training and testing were done using the rsfMRI data of 100 participants from the human connectome project (HCP), and the results establish the viability of the proposed model. Furthermore, experiments demonstrate that A-GHN outperforms the existing methods in learning the complex nature of human brain function.

## 1. Introduction

The human brain’s structural topology is estimated from diffusion tensor images (DTI) to derive the structural connectivity (SC) matrix that summarizes the fiber connectivity density among the brain regions. On the other hand, static or steady-state functional connectivity (FC) among these regions is estimated by computing the correlation coefficient (usually Pearson) of the respective timevarying resting-state functional magnetic resonance imaging (rsfMRI) signals. The correlation captures the spontaneous brain activity when participants are not engaged in any specified task. Furthermore, the brain activity observed in the rsfMRI signals is constrained and influenced by the connectome [1, 2]. Characterizing the SC-FC mapping is an open and challenging research problem in cognitive neuroscience [3]. Such models can be used to identify the biomarkers that underlie any deviation from the expected FC based on the SC in various diseases such as Autism Spectrum Disorder (ASD), Dementia, and many more [4, 5]. These models will also be helpful to characterize the functional recovery patterns resulting from therapy by comparing the FC observed with the predicted FC based on healthy structural topology [4, 5].

Traditionally, graph-based modeling has been popular for solving SC-FC mapping. One of the seminal works in this direction by [3] formulated a linear model that considers diffusion of regional brain activity over the graph topology by choosing a single optimal diffusion kernel. Later, [6, 7] utilized multiple diffusion kernels for learning the SC-FC mapping and demonstrated the superiority of using multiple diffusion kernels. The idea of multiple diffusion kernels formulation is specifically interesting and demonstrates the integration of multiple kernels within the same machine learning model. Becker et al. [8] proposed a versatile nonlinear mapping approach to obtain the functional connectivity from the structural connectivity random walks using spectral graph theory. In another relevant work on functional brain connectivity, [5] derived a relation between SC and FC via Laplacian spectra, where FC and SC share eigenvectors and their eigenvalues are exponentially related. However, most of these methods suffer from the computational overhead related to scalability or exhibit sub-optimal performance on SC-FC mapping over brain graphs. In another relevant work on functional brain connectivity, Abdelnour et al. [9] derived a relation between SC and FC via Laplacian spectra, where FC and SC share eigenvectors and their eigenvalues are exponentially related. However, most of these methods suffer from the computational overhead related to scalability or exhibit sub-optimal performance on SC-FC mapping over brain graphs.

A novel deep learning method called graph convolution network(GCN) [10] has recently been proposed to generalise convolutional neural network models for graph data.GCNs achieve state-of-the-art results in various application domains such as, computer vision [11], applied chemistry [12], natural language processing [13], and neuroscience [14]. The GCN-based Encoder-Decoder network was proposed for SC-FC mapping where the normalized Laplacian of SC was provided as input to GCN, and training was accomplished using the ground truth FC with an MSE loss function [15]. The primary limitation of this method is the absence of diffusion over multiple scales that is useful for the integration of information from the node attributes and the network topology. A recent variant of GCN, *GraphHeat Network* (GHN), attempts semi-supervised classification [16] and enables control over heat diffusion scales while filtering out the influence of high-frequency spectral components of the graph Laplacian. Another recent work proposed by [17] learned the graph kernels based on the intuition of the specific application domain instead of choosing standard kernels such as heat kernels and normalized heat kernels. However, the reported performance is poor on the SC-FC mapping experiments. Deep learning models such as CNNs and LSTMs, including GCNs, require large datasets for training and evaluation. However, recent benchmarking frameworks demonstrate how to work with smaller datasets for graph neural networks. They advocate the use of baselines to compare the performance of the proposed model using hyperparameter tuning and several validation experiments when using smaller graph datasets [18]. Hence, in this paper, we evaluated our A-GHN model on the smaller dataset (i.e 100 SC-FC pairs) to learn the complex FC structure.

Recently, attention mechanisms have become popular and standard to enable working with variable size inputs and for focusing on the most relevant parts of the input to make decisions [19, 20]. In the proposed model, A-GHN incorporates the propagation and aggregation of node representations by heat diffusion mechanism at multiple scales over the SC matrices. It is expected that the multiple scales contribute differently to the predicted FC. Thus, we introduce the attention mechanism to capture the contribution of each scale-specific A-GHN sub-models for learning the SC-FC mapping.

In summary, these methods together establish the usefulness of single or multi-scale diffusion [3, 7] and the feasibility of graph neural networks for solving the SC-FC mapping problem [15]. Inspired by these and the attention mechanism over multiple scale-specific GHNs, we propose an attention-based fusion of multiple GraphHeat networks (A-GHN) that efficiently employs multi-scale diffusion to promote computational tractability and scalability. Figure 1 display the pipeline of A-GHN model. The A-GHN model utilizes multiple GHNs, each with an independent channel of input based on heat kernels. Predicted FC is then computed based on the weighted combination of these outputs and is compared with the empirical FC. Here, the attention scores are computed by taking the softmax over weight coefficients, where each attention score corresponds to the A-GHN sub-models output. As a result, the proposed model approximates the empirical FC well, and the FCs recovered with the A-GHN approach seem to have better correspondence with the ground truth than related models that incorporate either multi-scale diffusion or GHN.

**Figure 1:**
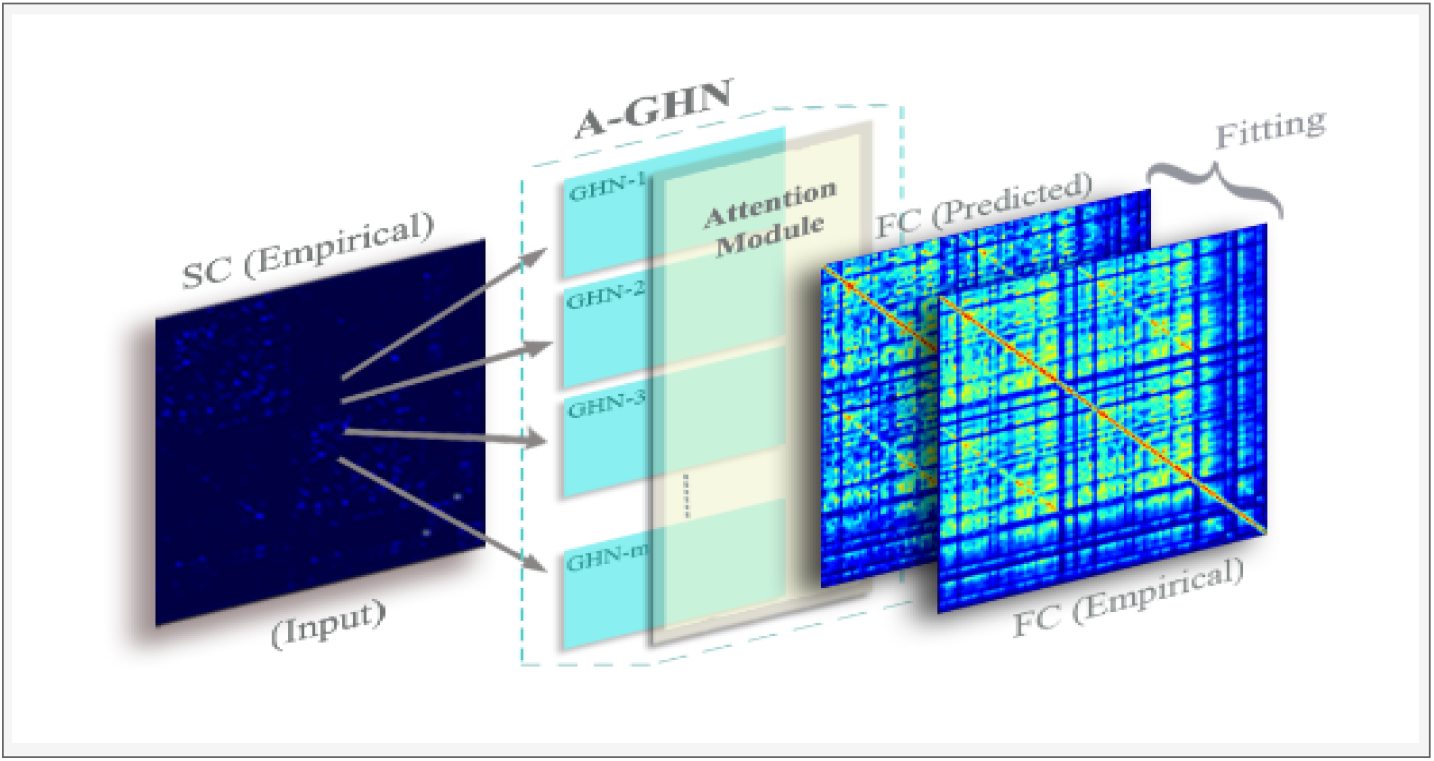
Mapping the structural and functional connectivity in brain graphs using the proposed A-GHN network.

The key contributions of this paper are the following:

- We propose a novel, end-to-end learnable A-GHN architecture for learning the SC-FC mapping on brain graphs.
- Our method is grounded in the theory of the reaction-diffusion process in the cognitive domain while retaining the key properties of generalisability, scalability, and tractability in the deep learning framework.
- We present a comprehensive empirical analysis including perturbation experiments and a detailed ablation study to demonstrate the proposed model’s robustness and validity on a publicly available dataset.

## 2. Related Work

### 2.1. Whole Brain Modeling of SC-FC

Classical methods proposed non-linear models of cortical activity, which were then extended to model whole-brain behavior via coupling between regions based on structural connectivity [21]. Also, the whole-brain computational models have been used as powerful tools to understand the relationship between structural and functional brain connectivity by linking the brain function with its physiological underpinnings [22, 23, 24]. Several other studies place non-linear oscillators at each cortical location and likewise couple them using anatomic connectivity strength [25, 26, 27]. However, these simulation models are only revealed through large scale, fine-grained stochastic simulations over thousands of time samples, and pose a practical challenge for the task of inferring functional connectivity from structural connectivity.

### 2.2. Graph-theoretic Modeling using Linear Models

The earlier graph-theoretic modeling experiments studied the mapping of SC-FC relationships by capturing the correlation structure of whole-brain dynamics using linear models [3]. Specifically, [3] present a simple, low-dimensional network diffusion linear model producing an accurate description of the SC-FC relationship. However, this model uses one global parameter across all the subjects, and the hypothesis of a single scale best-fitting kernel across subjects is not tenable. Surampudi et al. [6] observed that the combination of multiple diffusion scales exhibits scale-dependent relationships among various regions of interest (ROIs), and these multi-scale diffusion kernels can capture reaction-diffusion systems operating on a fixed underlying connectome (SC). However, multiple diffusion kernels were not sufficient to explain the self-organizing resting-state patterns found in FC. Recently, a new framework, the multiple kernel learning model (MKL), provides plausible mathematical reasoning for the existence of these co-activations along with diffusion kernels by linearizing a variant of the reaction-diffusion model and extending it to generate FC [7].

### 2.3. Deep Learning Models for SC-FC Mapping

Recently, the study of GCNs has successfully reconstructed the brain FC from SC graph by building a graph encoder-decoder system [15]. Moreover, the learned low-dimensional embeddings capture essential information regarding the relationship between functional and structural networks. However, the major limitation of this method is the lack of control over a single diffusion kernel at an optimum scale and multiple scales of diffusion. On the other hand, [17] proposed a deep graph spectral evaluation network (GSEN) for modeling the graph topology evolution by the composition of a newly generalised kernel. This method efficiently models the global and local evolution patterns between the source (SC) and target (FC) graphs. Although the method seems interesting, the GSEN model reports a poor performance on SC-FC mapping.

## 3. Proposed Solution

### 3.1. Problem Statement and Proposed Solution

The brain is typically represented as a graph in the computational neuroscience community, where graph nodes are modeled as key brain regions, and edges represent their structural or functional relationships. The aim here is to learn a mapping between the two brain graphs representing a sparse structural connectivity matrix (SC) and a dense static (steady-state) functional connectivity (FC) matrix, as depicted in Figure 2. We propose to employ multi-scale heat diffusion kernels in a novel deep learning framework for this task.

**Figure 2:**
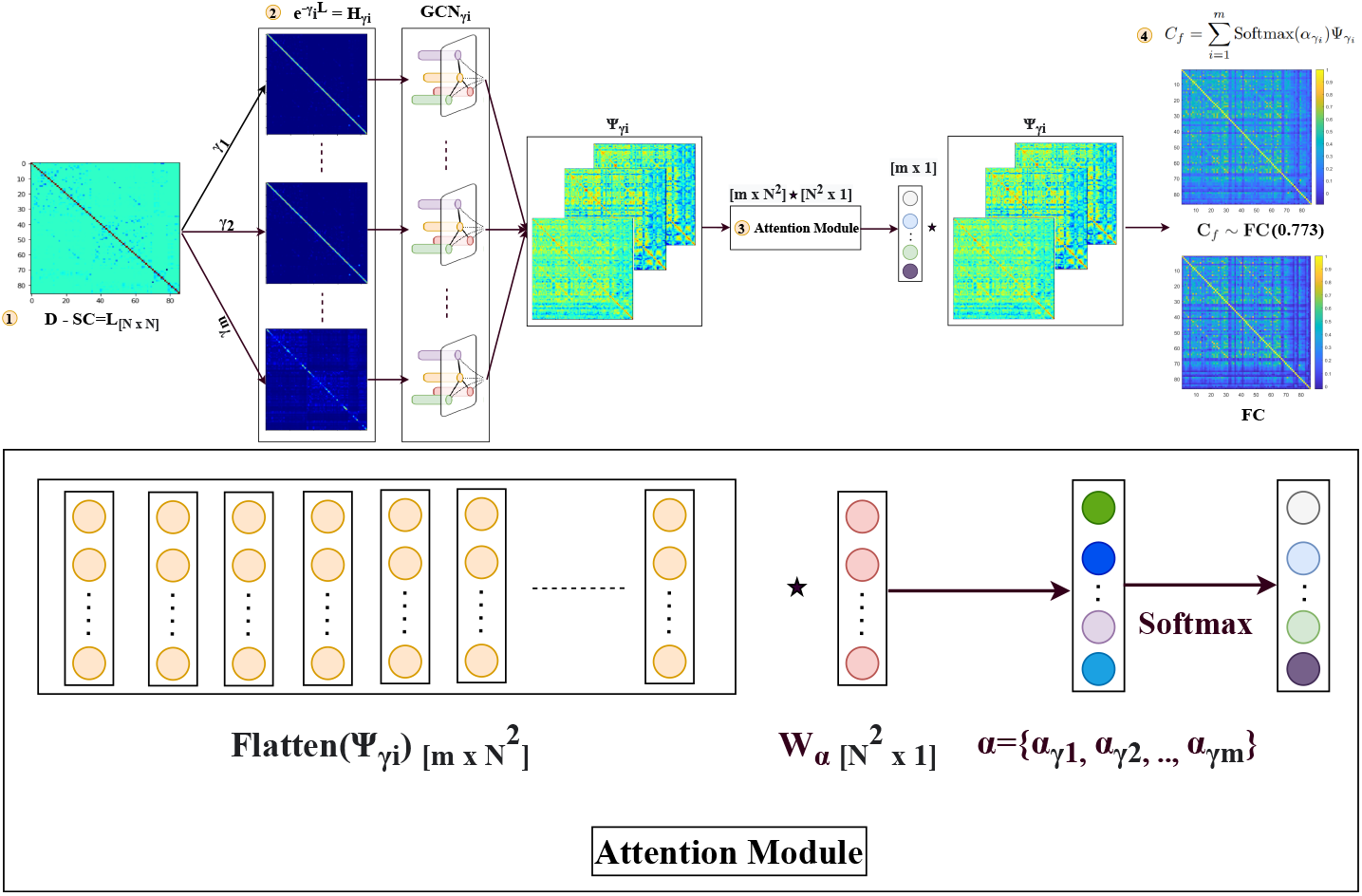
Proposed A-GHN architecture for learning SC-FC mapping using multi-scale GraphHeat networks (GHN) along with attention mechanism. A Laplacian matrix is computed from the structural connectivity matrix (SC) input in step 1. Multiple heat kernel matrices are obtained using m different diffusion scales and fed to the individual (A-GHN sub-model) in step 2. In step 3, an attention module is introduced to learn the attention scores corresponding to A-GHN sub-models. A Softmax linear combination of the outputs Ψ_*γ_i_*_ yields the predicted functional connectivity (*C_f_*), which is compared with the ground truth empirical FC in step 4.

### 3.2. Mathematical Background & Notations

#### Graph Definition

Consider a weighted, undirected graph denoted by 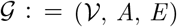, where 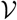 is a set of *N* nodes, 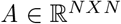 is the symmetric adjacency matrix and *E* is the set of edges connecting the nodes. A graph Laplacian matrix is defined as *L* = *D* – *A*, where *D* is a diagonal matrix with degree of nodes on the diagonal, 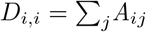 The spectral decomposition of the Laplacian matrix (*L* = *U*Λ*U^T^*) yields (i) Eigenvector matrix (*U*) and (ii) Eigenvalue matrix (Λ) which is a diagonal matrix with the eigenvalues arranged in increasing order.

#### Graph Convolutional Networks

Graph Convolutional Neural network (GCN) is a multi-layer neural network that convolves neighboring node’s features and propagates a node’s embedding vectors to its nearest neighborhood [10]. For a one-layer GCN with Z hidden units, the latent node feature representation (Ψ^(1)^) is computed as

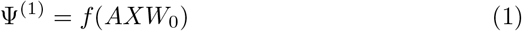

where *A* is the symmetric adjacency matrix, 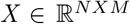 is the node feature matrix where each row of the matrix represents a M-dimensional content vector for each node in the graph, 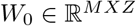 is weight parameter associated with the 1^*st*^ layer of GCN, and *f* is activation function. One can incorporate higher-order information of the neighborhoods by stacking multiple GCN layers

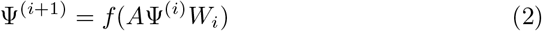

where *i* denotes layer number and Ψ^0^ = *X*

#### Graph Convolution using Heat Kernel

The *GraphHeat Network* (GHN) formulation captures the smoothness of labels or features over the neighborhood of the nodes as determined by the graph structure [16]. A heat kernel is defined as

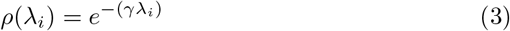

where *γ* ≥ 0 is the scale hyper-parameter, and *λ_i_* denotes the *i^th^* eigenvalue in Λ. Let 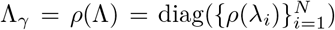 denote the kernelized diagonal matrix. Thus, we can define the convolution kernel (*g_w_*) as

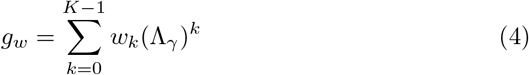

where *w_k_* is the weight parameter and here we choose *K* = 2 (*i.e*. only considering the first-order polynomial approximation of ChebyNet [28]).

For the given input signal *X*, graph convolution is achieved as follows:

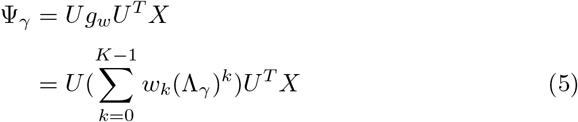

Specifically, for our choice of *K* = 2:

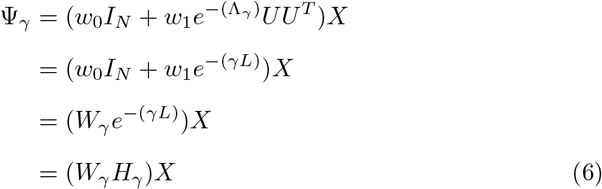

where 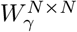 is a weight matrix corresponding to scale *γ, H_γ_* = *e*^−*γL*^ represents the heat kernel matrix, and 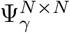 is the scale-specific output of GHN.

### 3.3. Attention based Multiple GraphHeat Networks (A-GHN)

Let us consider *m* heat kernel matrices with *m* different scales {*γ*_1_, *γ*_2_, *γ*_3_, ⋯, *γ_m_*} and corresponding GraphHeat kernels {*H*_*γ*_1__, *H*_*γ*_2__, *H*_*γ*_3__,···, *H_γ_m__*}. A-GHN already includes the propagation and aggregation of node representations by heat diffusion mechanism over the SC matrix. Further, the weight matrix parameters associated with the structural graph are learned during the model training process, reflecting the mean regional activities. Hence, the node feature vector X was chosen as a one-hot vector (*I_N_*) in our model setting.

Each A-GHN sub-model outputs a matrix Ψ*_γ_i__* and we hypothesize that the linear combination of the softmax probabilities with A-GHN sub-model outputs would give rise to a good estimate of FC. Let *α* = {*α*_*γ*_1__, *α*_*γ*_2__, ⋯, *α_γ_m__*} denote the weight coefficients in the linear combination corresponding to the *m* GHN branches (A-GHN sub-models). These weight coefficients are learned by feeding the outputs of all *m* GHN branches to a fully connected layer. In our proposed A-GHN model, the attention module is designed such that the differential contribution of multiple scales is weighted appropriately to estimate the predicted FC. In order to obtain the normalized weights (attention scores), we utilize the softmax activation function. Finally, the linear combination of the outputs of *m* GHNs weighted by the corresponding attention scores allows us to jointly train all A-GHN sub-models and the fully connected layer via end-to-end back-propagation learning.

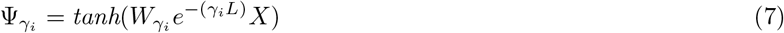

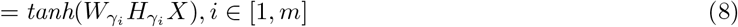

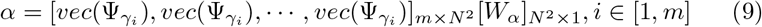

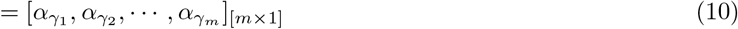

where *α_γ_i__* = *vec*(Ψ_*γ_i_*_) × *W_α_* denote the linear coefficients capturing contribution of the individual heat kernel Ψ*_γ_i__*.

Thus, we approximate the empirical FC with weighted combination of output of multiple A-GHN sub-models corresponding to *m* diffusion scales to predict the FC (*C_f_*) as follows

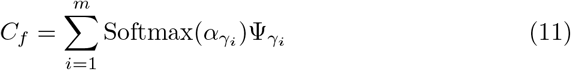

#### Loss Function

The attention parameters *W_α_* and scale-specific parameters *W_γ_i__* are estimated from the training subjects (indexed by *s* that varies from 1 to *S*) and remain fixed during the testing phase. We consider the loss function *J* (Equation of 12) to be the mean squared error between empirical and predicted FCs. Since the target FC matrix is symmetric, we have also made the estimated FC matrix (*C_f_*) symmetric by adding its transpose, similar to MKL [7]. The loss function is then minimized using the stochastic gradient descent procedure.

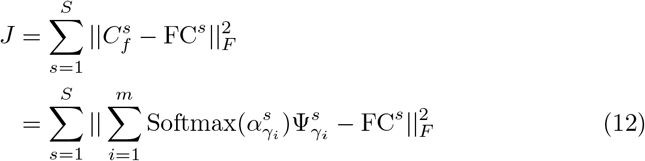

Here, 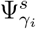 denotes a *N* × *N* matrix with subject index (s) and *α* denotes an attention *m* × 1 vector. Figure 2 depicts the proposed architecture that combines attention-based fusion of A-GHN sub-models with multiple heat kernels.

### 3.4. Relation to Reaction Diffusion Phenomenon

Mutual interaction of the elements of a complex system results in a neural field of activity which in turn leads to the formation of self-organizing patterns. Reaction-Diffusion (RD) model is the mathematical framework that characterizes such a spatio-temporal change in the field. RD systems have been successfully used to model the interaction among neurons belonging to different brain regions and the associated functional connectivity (FC) among the regions of interest (ROIs) of the brain [29, 30]. The reaction part of the RD model corresponds to the interaction of the excitatory and inhibitory neural elements, and the diffusion part corresponds to the spreading of the resultant neural activity over the structural fiber pathways. As the interacting (reacting) neural elements differ in their parameters, the emerging spontaneous activity of the neural ensemble results in non-linear patterns. The growth and the progression of a neural field are mathematically characterized by the Wilson-Cowan model, a variant of the RD framework. The statistical behavior of the mean activity of the neural fields is described by the equations of the Wilson-Cowan model [31, 32].

Inspired from the multiple kernel learning model (MKL) model [7] which is based on the RD framework, in this paper, we propose attention-based multiple GraphHeat networks (A-GHN) to map SC-FC. The proposed solution formulation is analogous to MKL and is as follows:

#### MKL

The optimization formulation minimizes an objective function *J* comprising the mean squared error between empirical and predicted FCs as in [7] and is represented as:

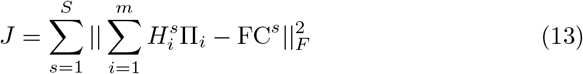

where, II_*i*_ are estimated from the training subjects (indexed by *s* that varies from 1 to *S*), and 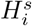, denotes the Heat Kernel matrix of subject s associated with scale *i*.

Similarly, in [6], the mixing coefficients are subsequently learned while solving an optimization formulation as:

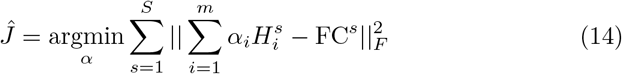

where *α_i_*, is a weight coefficient associated with scale specific heat kernel *H_i_*

From Equations 7, 12, and 13, we observe that the learnable parameters 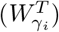 in Equation 12 in the proposed framework are analogous to the estimated parameters (Π_*i*_) in Equation 13 of the MKL framework [7]. Thus, as hypothesized in [7], we can interpret 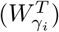 as corresponding to the initial mean regional activities. Hence, Ψ_*γ_i_*_. in Equation 12 of the proposed framework, when viewed along with Equation 7, would correspond to the diffused output based on the initial mean regional activities.

Additionally, we introduce an attention mechanism in our proposed model (A-GHN) that combines attention scores with the outputs of *m* GHNs. From Equations 12, 14, the learnable mixing coefficients through optimization formulation in Equation 14 are analogous to the weighted attention scores obtained through gradient descent in Equation 12.

We present the visualizations of (*W_γ_i__*) and the correlation plot between the empirical and predicted FCs without attention in section 4.5.

## 4. Experimental Setup & Results

This section provides details of the experimental setup, dataset, model design, and comprehensive evaluation of the proposed model. Further, we performed detailed ablation studies where we induced perturbations in the input and conducted studies by removing the attention module to see the impact on the performance in all the cases and justify the proposed architecture.

### 4.1. Dataset

Deep learning models typically require a large amount of data for training as they involve the learning of a huge number of parameters. Further, MRI data acquisition comprising different modalities such as T1, DTI, and rsfMRI is a costly and time-consuming process. In light of these issues and in order to obtain a meaningful comparison against the existing results, we considered a popular and widely used dataset from the human connectome project (HCP). We have considered the structural connectivity - functional connectivity (SC-FC) pairs of a total of 100 subjects from the HCP repository (see [33] for data pre-processing methodology). All these participants underwent resting-state functional imaging (no task condition) with their eyes closed. The blood oxygen level dependent (BOLD) time-series signal available for each participant has 1200 time points aggregated across 86 regions of interest (ROIs) as per the AAL brain atlas [34]. We omitted data of 2 participants because of quality issues in the pre-processing stage; hence data of 98 participants were considered for further experiments.

### 4.2. Baseline Methods

Since the proposed model combines graph convolutional network with multiple heat kernel diffusion, we chose two related baseline methods for comparative analysis. The first method, multiple kernel learning (MKL) model proposed in [7], utilizes multi-scale diffusion over brain graphs to learn the subject’s SC-FC mapping but does not incorporate deep networks. On the other hand, the second method uses GCN-based Encoder-Decoder architecture [15] is a deep learning-based model. However, this does not incorporate multi-scale diffusion. Thus, the two baselines together allow us to evaluate the impact of deep networks and that of the multi-scale diffusion independently against our proposed A-GHN model. We replicated both the MKL and GCN Encoder-Decoder models with the same choice of parameters as indicated in the original papers on the data from 98 participants from HCP for training and testing experiments.

### 4.3. Model Setup

Here, we describe the model setup, training and testing phases for the proposed A-GHN model.

#### Training Phase

We trained the A-GHN model on HCP rsfMRI data where a randomly chosen set of 45 subjects was used for training (45 SC-FC pairs), 5 subjects (5 SC-FC pairs) for validation and the remaining 48 subjects (48 SC-FC pairs) for testing. The 86 × 86 heat kernel matrix obtained from the Laplacian of structural connectivity (SC) matrix was given as input to the graph convolution networks (GCN) and the 86 × 86 empirical functional connectivity (FC) matrix as the target output to train the model. Here, the number of vertices corresponds to the 86 brain regions, and the edges represent the structural fibers connecting the brain regions over which heat diffusion takes place. As shown in Figure 2, outputs of the one-layer A-GHN models were combined in a weighted manner using the corresponding attention scores obtained from the softmax layer. The number of coefficients obtained is equal to the number of scales (*m*=7), and the final output is an (86 × 86) predicted FC. We used mean squared error (MSE) between empirical and predicted FC matrices as the loss function for learning.

#### A-GHN hyper-parameters

To perform SC-FC mapping using A-GHN, we set the convolution layer’s embedding size as 86 and the input node feature vector *X* as the identity matrix (*I_N_*). We used Adam optimizer [35] with an initial learning rate of 0.001, *tanh* as the activation function, and the *L*_2_ weight decay was set to 5*e*^−4^. We applied dropout with a keep-probability of 0.5 and trained the A-GHN model for a maximum of 100 epochs. To overcome the overfitting problem, we stopped training if the validation loss did not decrease for 10 consecutive epochs (See supplementary material for the profiles of learning curves in Figure S1).

#### Testing Phase

We used the other half (48 SC-FC pairs) to predict the corresponding FC matrices in model testing. We followed the same parameters used in model training except omitting the drop-out parameter. We use the Pearson correlation coefficient between the empirical (ground truth) and the predicted functional connectivity (FC) matrices to measure the model performance. There were three kinds of validation experiments performed – 5-runs (each run with random initialization), 5-fold cross-validation (CV), and Leave-one-out crossvalidation (LOOCV) experiments. For the 5-runs set-up, we have a train-test split of data of 50 and 48 participants. While the test set of 48 SC-FC pairs remains the same for all the experiments, in each of the five runs, a random set of 45 participants is chosen for training and the remaining 5 participants for validation in order to avoid any bias in the training and validation sets. Five such runs were performed, and the average results for each test subject are shown in Figure 4. For the 5-fold CV experiments, 4-folds are used for training and one-fold for testing. The results of the 5-fold CV are shown in Figure 6. LOOCV is conducted in a regular manner where the train-test splits comprise 97:1 participants. Of the 97 participants’ data, 90 were used for training and 7 for validation. For ease of comparison, we depict the results of 48 test participants in Figure 7 with the same index numbers as in the 5-run experiments shown in Figure 4. The LOOCV results of all the 98 participants are shown in Figure S4 (in the supplementary). Thus, the validation results establish the generalisability of the results with different data splits.

#### Choice of Model Parameters

The choice of various model parameters is explained below.

#### Choice of *m*

Figure 3 shows the profile of heat kernels for various scales of diffusion ( γ) ranging from 0.5-10. The GraphHeat formalism [16] allows for selective focus on low-frequency spectral components at higher scales, whereas high-frequency spectral components are suppressed at lower scales. Hence, in this paper, we chose multiple scales where each scale of diffusion characterise to determine neighboring nodes that reflect the local structure or the relevant information of smoothness manifested in the graph structure. As can be seen in Figure 3, the local diffusion phenomenon is observed for smaller scales (0.5-1) with contribution from many eigenvalues/vectors, including the large eigenvalues. On the other hand, the global diffusion phenomenon is noticed for bigger scales (1-10) that depend predominantly on the contribution from eigenvalues/vectors corresponding to smaller eigenvalues. The number of heat diffusion scales (see Eq. 7) was set to *m* = 7 empirically, based on the performance of the proposed model. We used ascending order of scales that correspond to the global diffusion phenomenon in case of lower scale indices (*γ* values of 0.6 and 0.8) and local diffusion phenomenon in case of higher scale indices (*γ* values of 1, 2, 4, 6, and 8) (see Figure 3).

**Figure 3:**
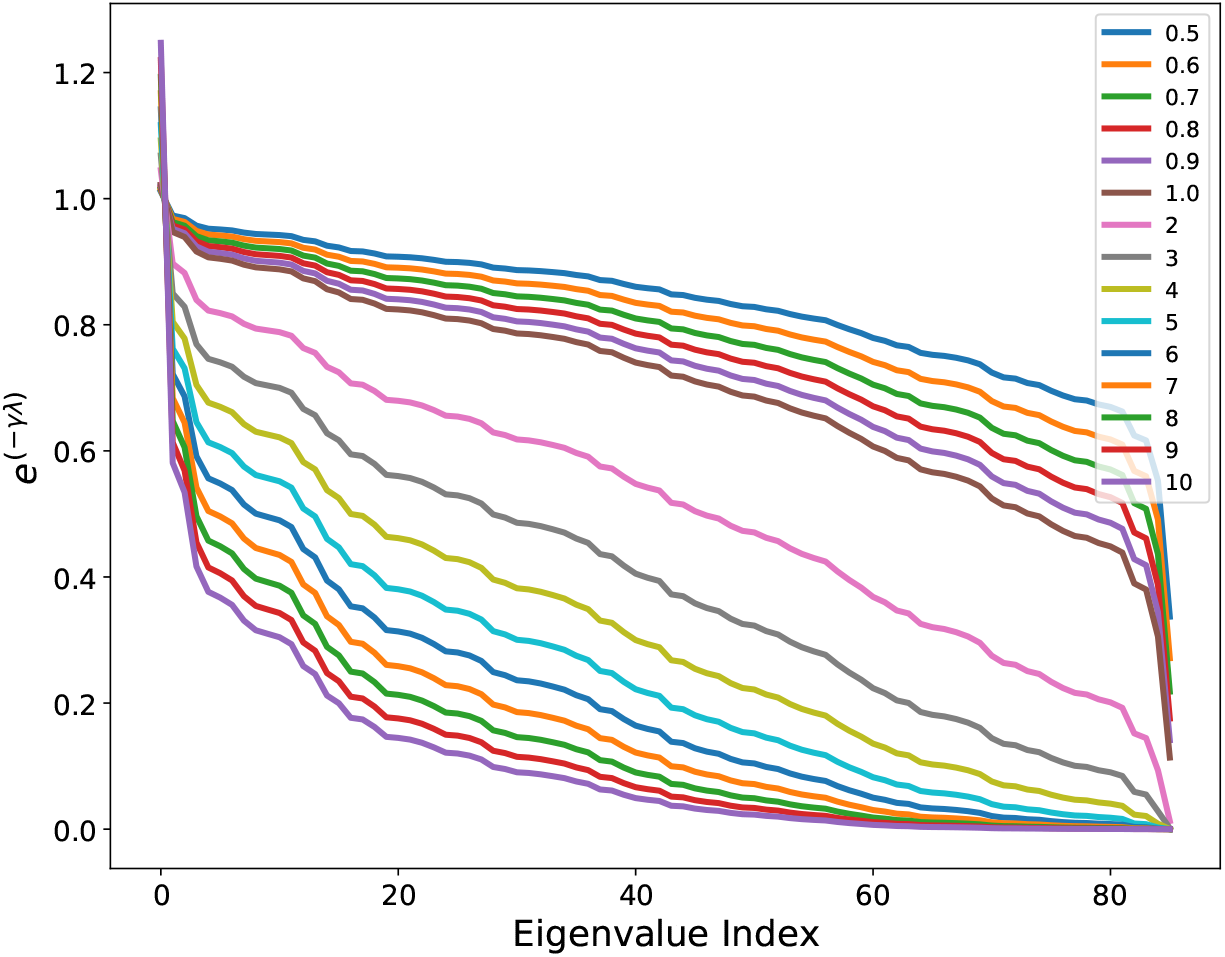
Depicts different diffusion scales (*γ*) ranging from 0.5-10 (values in the legend), and each exponential curve is a function of the scale (*γ*) and represents the contribution of every eigenvalue of the Laplacian of the SC matrix (the indices of eigenvalues (in increasing order) are shown on the abscissa).

#### Choice of Activation Function

In order to determine the kind of activation function to be used in the output layer, we ran experiments with several choices and found that *tanh* is suitable. We observed that *tanh*, *relu*, and *leaky relu* (with a negative slope of 0.01) activation functions yielded similar performance values while the configuration with *sigmoid* function had a lower performance. Since the *tanh*-version showed a slight edge over the others, *tanh* was chosen as the activation function in the output layer of the A-GHN for further experiments. These results are shown in Figure S2 in the supplementary material.

#### Choice of A-GHN Layers

To understand the impact of increasing the number of hidden layers of A-GHN, we experimented with a two-layer A-GHN model. We empirically found that the mean Pearson correlation of test subjects with the two-layer model was lower than that of the one-layer model, as shown in Figure S3 (supplementary material). It appeared that an increase in the number of layers led to over-fitting and a decrease in performance. Hence, we considered a one-layer A-GHN model for all the experiments.

### 4.4. Results

#### Quantitative Evaluation

We compared the performance of our proposed model with two existing approaches: Multiple Kernel Learning (*MKL*) model [7] and the GCN-based Encoder-Decoder model [15]. The results of the comparative study using the 5-random-run experiments are shown in Figure 4, where we can see that the proposed A-GHN model performs better with a mean correlation value of 0.773 in the range of [0.60, 0.87] on the test set as compared to GCN-based Encoder-Decoder model (Mean = 0.73, range in [0.53, 0.82]) and MKL (Mean = 0.69, range in [0.27, 0.87]). In order to estimate the statistical significance of the performance differences, we performed One-way ANOVA on the mean correlation values for the test participants across the three models. The main effect of model was significant [F(2,141)=10.26, *p*=.00007]. Further, the *post hoc* pairwise tests revealed that the mean correlation values of the A-GHN model were significantly different from those of the other two models [with GCN Encoder-Decoder: *p*=.03 and with MKL: *p*=.00004]. On the other hand, the performance of the two baseline models did not differ significantly [GCN Encoder-Decoder vs. MKL: *p*=.12].

**Figure 4:**
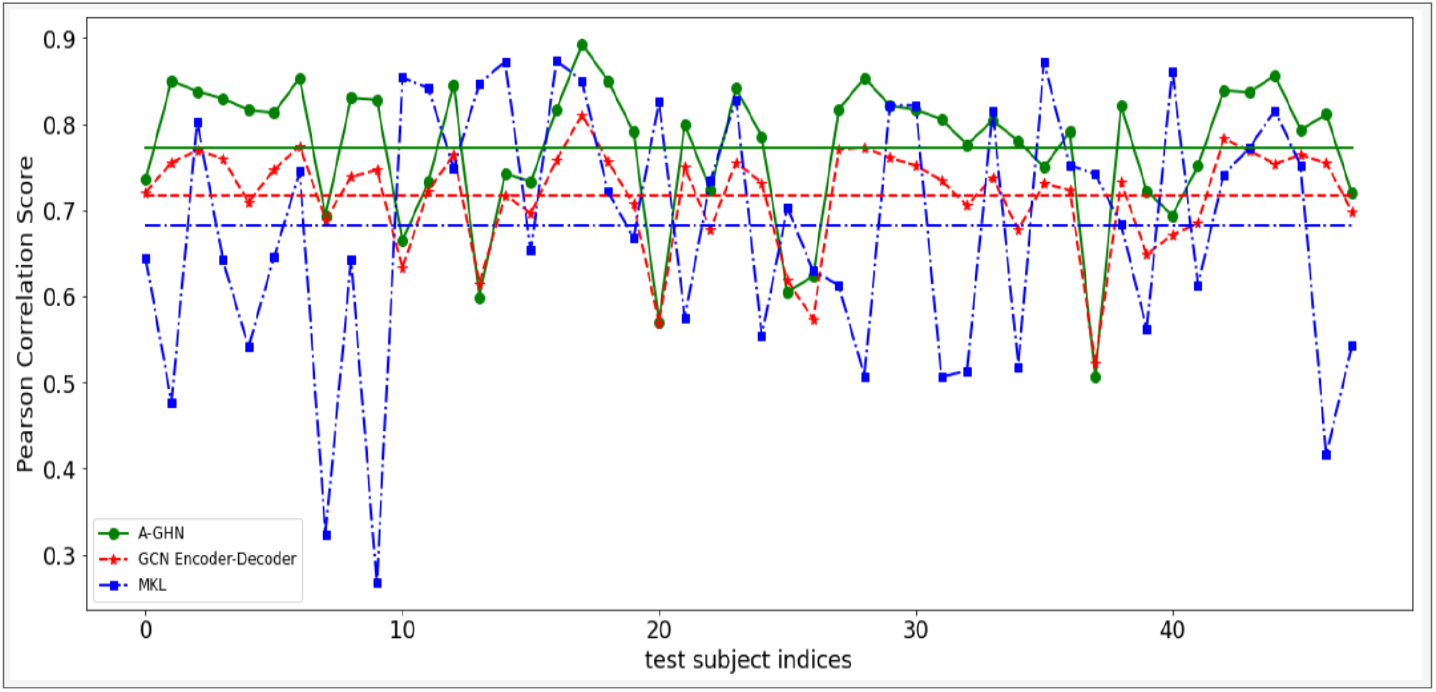
Pearson correlation values between empirical and predicted FCs of all the test subjects with the proposed A-GHN model (Green line), averaged over five runs, are compared with the predictions of the other two models. Horizontal lines show the mean correlation values (higher is better) of 0.78, 0.73, and 0.69, respectively, for A-GHN, GCN Encoder-Decoder, and MKL.

Similarly, Figure 5 display the mean squared error (MSE) of test subjects using the 5-random-run experiments, where the proposed A-GHN performs a lower MSE value of 0.0265 in the range of [0.013, 0.054] on the test set as compared to GCN-based Encoder-Decoder model (Mean = 0.037, range in [0.024, 0.067]) and MKL (Mean = 0.086, range in [0.015, 0.261]). Further, the statistical significance test using the one-way Anova test provides an F-statistic [F(2,141) = 37.33, *p* = 0] concludes that the model was significant. Also, the post-hoc Tukey-HSD test reported that the proposed A-GHN model was significantly different with two models [with GCN Encoder-Decoder: *p*=.016 and with MKL: *p*=.00001].

**Figure 5:**
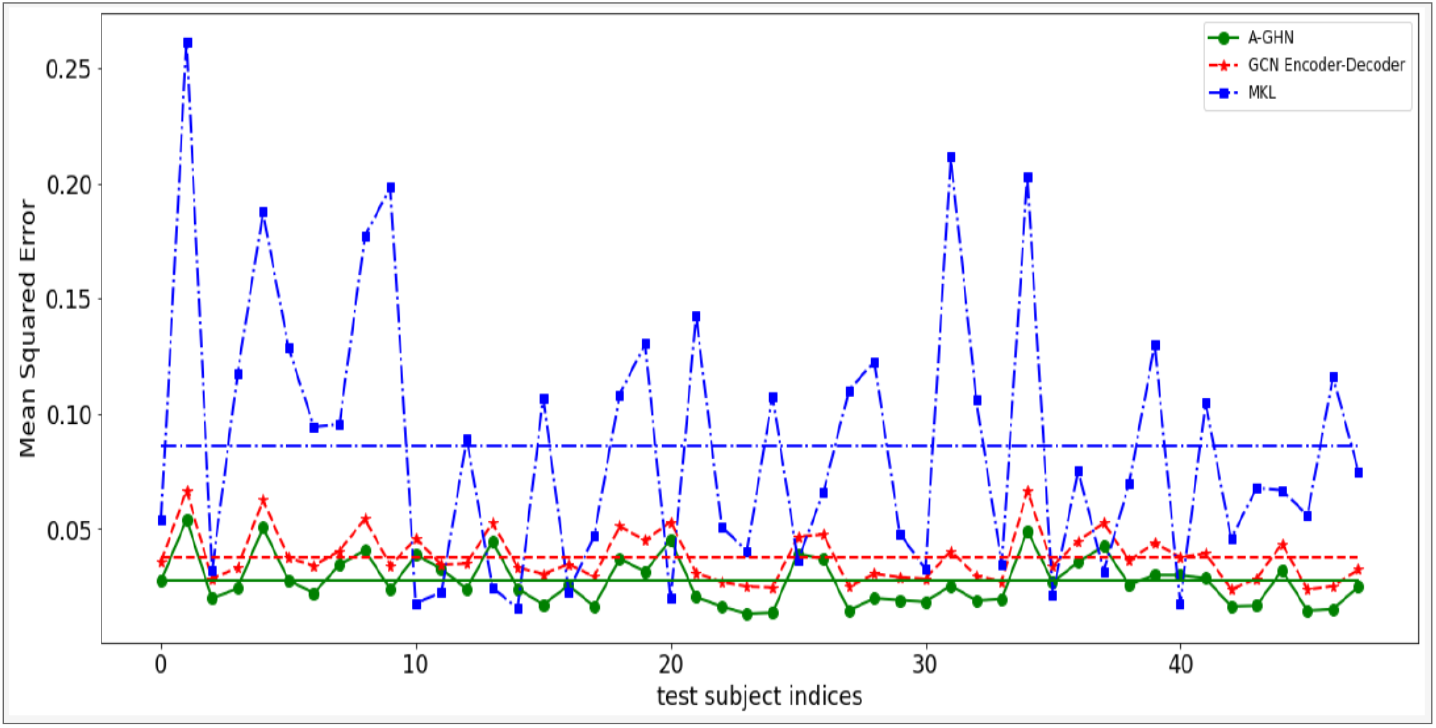
Mean Square Error (MSE) values between empirical and predicted FCs of all the test subjects with the proposed A-GHN model (Green line), averaged over five runs, are compared with the predictions of the other two models. Horizontal lines show the mean MSE values (lower is better) of 0.0265, 0.037, and 0.086, respectively, for A-GHN, GCN EncoderDecoder, and MKL.

**Figure 6:**
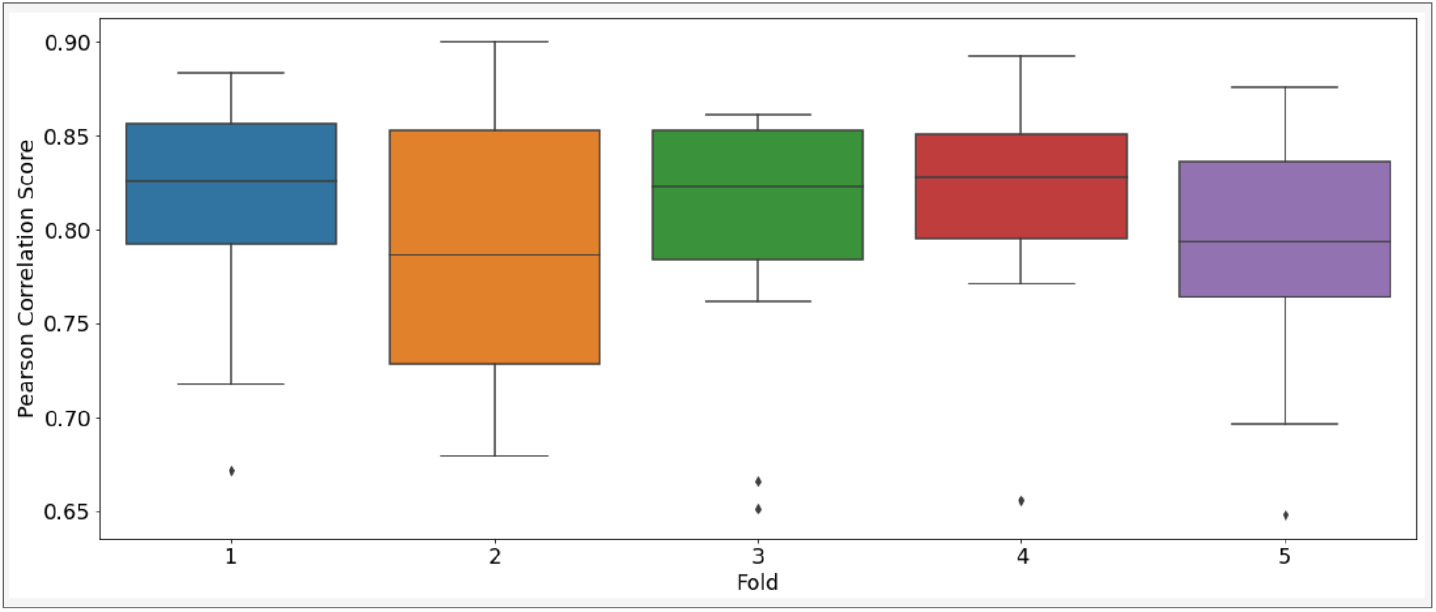
Results of performance of A-GHN model in the 5-fold cross-validation setting. The box plots depict the Pearson correlation between empirical and predicted FCs in each fold.

**Figure 7:**
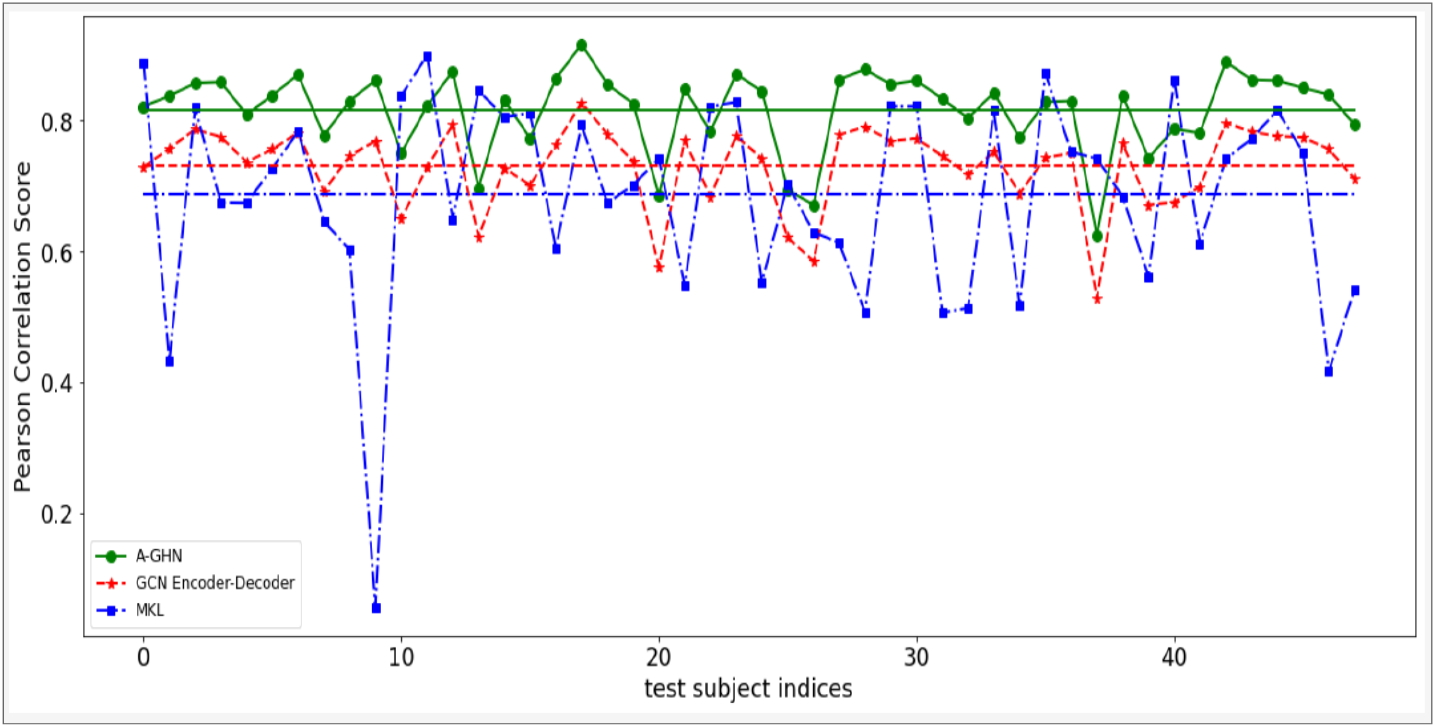
Pearson Correlation Results of leave-one-out cross-validation (LOO-CV) on the test subjects of A-GHN performs better than the other two models: GCN Encoder-Decoder and MKL. Note that the subject indices are kept identical to those in Figure 4.

##### Model Validation

To validate our model’s robustness, we performed various experiments such as leave-one-out cross-validation and 5-fold cross-validation to assess whether our model overfits the training data.

#### Qualitative Evaluation

We computed the mean of the predicted FC and the mean of the empirical FC matrices of the test subjects. We also computed the mean predicted FC matrices of the baseline models (GCN Encoder-Decoder and MKL). The visualizations of FC matrices are shown in Figure 8. Here, we can observe a better qualitative match between the mean predicted FC of our proposed model and the mean ground truth.

**Figure 8:**
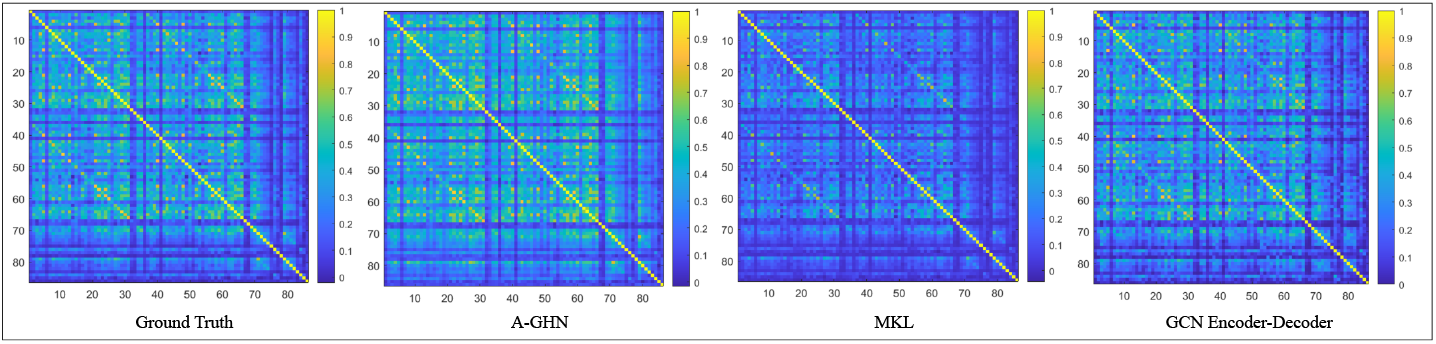
Qualitative comparison of the Functional Connectivity matrices (FCs). The mean of the predicted FCs from the proposed A-GHN model is compared with that of the mean FC from ground truth (empirically observed), GCN Encode-Decoder [15] and MKL [7] models.

In order to look at the finer details of the goodness of the learned mapping, four FC Networks were derived from the mean FC matrices of the test subjects using the Louvain algorithm available in the brain-connectivity-toolbox [36]. The edge-connectivity patterns of the predictions of the three models and the ground truth were rendered on a brain surface using BrainNet viewer [37] to understand the similarity of node and edge distributions between the empirical and the predicted FCs, shown in Figure 9. It can be seen that the proposed A-GHN model has a higher visual similarity to the empirical FC in terms of community assignment and inter-hemispheric connections as compared to the other models.

**Figure 9:**
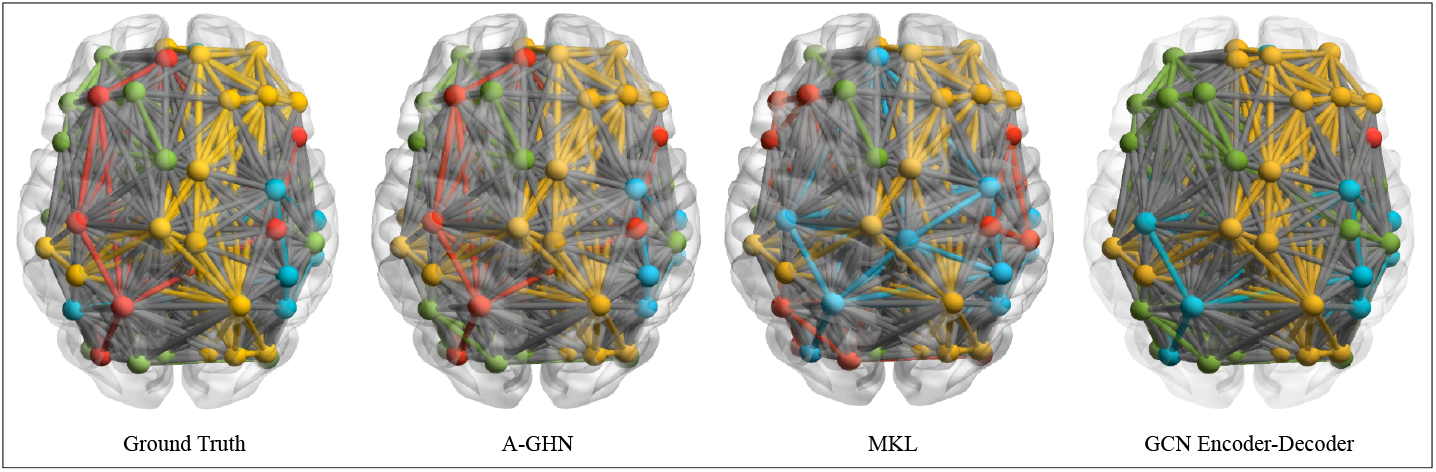
Qualitative comparison of the Functional Connectivity Networks. Four communities are derived from the mean FC matrices of the test subjects from the ground truth as well as the predicted FCs from the proposed and other models: MKL [7] and GCN EncoderDecoder [15]. Color coding of the edges/nodes for different models is done independently, and hence the cross-comparison of community structures is qualitative in nature.

### 4.5. Ablation Studies

We performed various ablation studies to establish the robustness of the proposed model. An ablation study was carried out to measure the importance of the attention module incorporated in the proposed model. To further verify whether our model learns the SC-FC relationship correctly and does not simply over-fit the data, we conducted two perturbation studies. One experiment studies the impact of perturbing the test input when the training protocol is kept intact. The second one verifies the results when the model was trained using perturbed inputs but tested on the original target outputs.

#### Importance of Attention

The distinguishing feature of the proposed A-GHN model is the use of attention in order to estimate a weighted combination of the GHN outputs. In order to assess the importance of the attention module, we performed an ablation study. The model was run without attention (called U-GHN in Figure 10) weights by simply summing and averaging the outputs of the seven A-GHN sub-models to obtain the predicted FC. It can be observed in Figure 10 that attention makes a difference in that the mean correlation value of A-GHN is 0.773 [range: (0.60, 0.87)] as compared to 0.741 [range: (0.46, 0.87)] of U-GHN. An F-test establishes that these differences are statistically significant [F(1,94)=5.3427, *p*=.023]. Similarly, we report the mean squared error (MSE) of test subjects using both A-GHN and U-GHN models in Figure 11. From Figure 11, we can observe that the overall MSE value of A-GHN is 0.0254 low as compared to 0.0287 of U-GHN.

To investigate the importance of attention-based A-GHN model vs. independently trained scale-specific models, we report the MSE values of individual scale-specific and A-GHN sub-models in Table 1.

**Figure 10:**
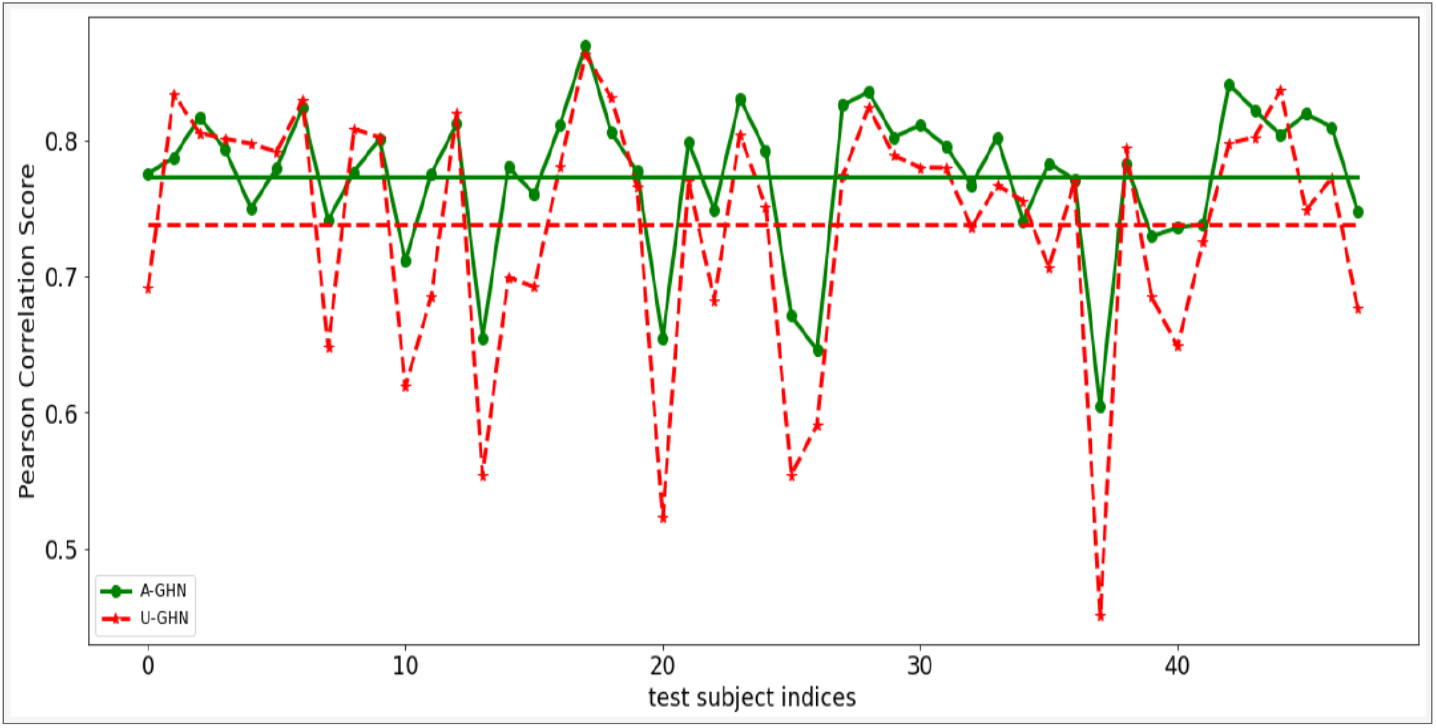
Results of Pearson Correlation (higher the better) comparison between A-GHN (with Attention) vs. U-GHN (Uniform GHN, without Attention) on the test subjects yield a comparative performance. Note that the subject indices are kept identical to those in Figure 4.

**Figure 11:**
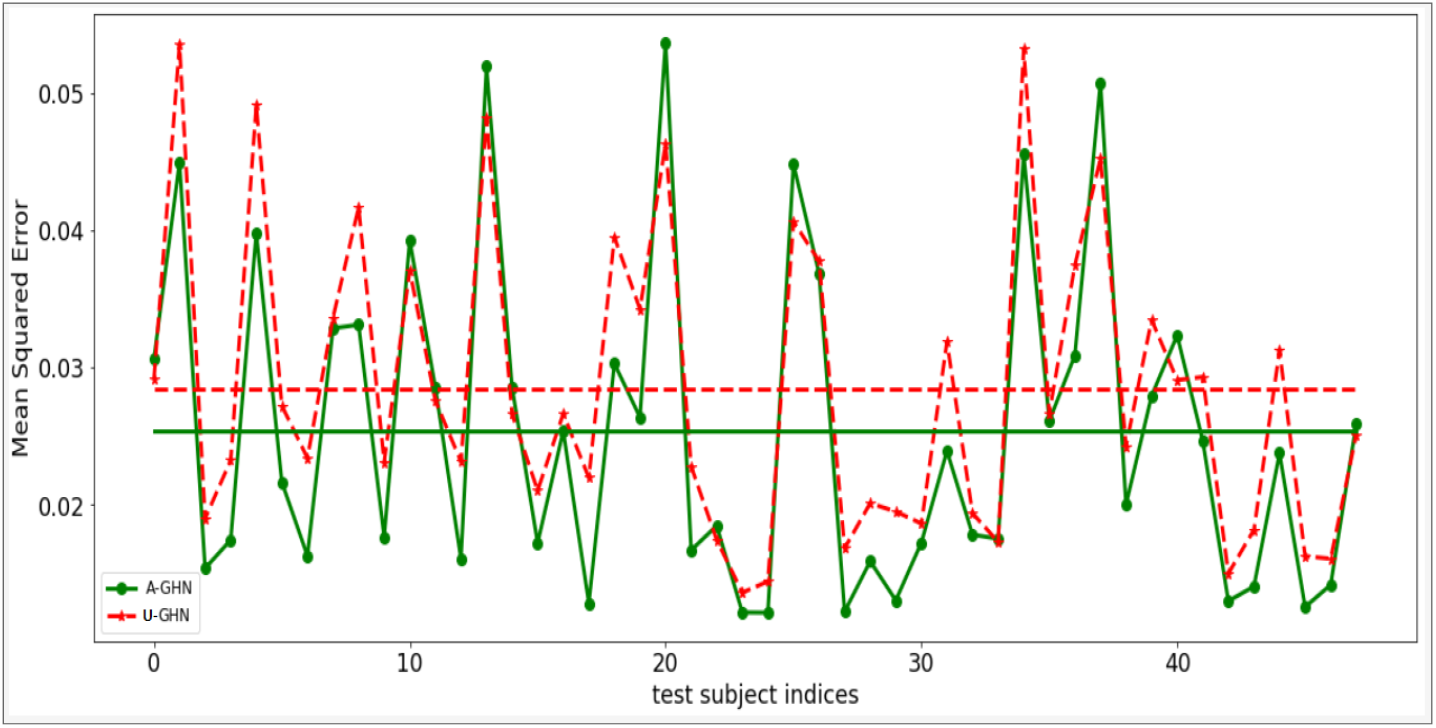
Results of MSE (lower the better) comparison between A-GHN (with Attention) vs. U-GHN (Uniform GHN, without Attention) on the test subjects yield a comparative performance. Note that the subject indices are kept identical to those in Figure 4.

**Table 1:** Comparison of A-GHN sub-models and individual scale-specific trained models of testing subjects. Comparison is done by computing the MSE between the ground-truth FC and predicted FC of test subjects. Overall, the A-GHN model displays a lower MSE value of 0.025 better than individual scale-specific trained models.

| Model Type→ | Independent Trained Model | A-GHN Sub-Model |
| --- | --- | --- |
| Scales | MSE | MSE |
| 0.6 | 0.0341 | 0.02840 |
| 0.8 | 0.0345 | 0.02841 |
| 1 | 0.0351 | 0.02845 |
| 2 | 0.0372 | 0.02940 |
| 4 | 0.0409 | 0.03271 |
| 6 | 0.0434 | 0.03533 |
| 8 | 0.0451 | 0.03662 |

#### Varying the Training Data Size

Figure 12 shows the mean Pearson correlations of the A-GHN model with varying training data set sizes. We ran the model with three different settings - 25%, 50%, and 75% of the data for training and subsequently tested with the remaining data. As expected, the mean correlation of the proposed model increases with the size of the training set – from 0.761 (25%), 0.773 (50%) to 0.779 (75%). However, the increase in performance is marginal as the A-GHN model yields a similar level of performance with a smaller training dataset, possibly because the graph convolutional network captures essential characteristics of the SC-FC mapping even with a sparse training set.

**Figure 12:**
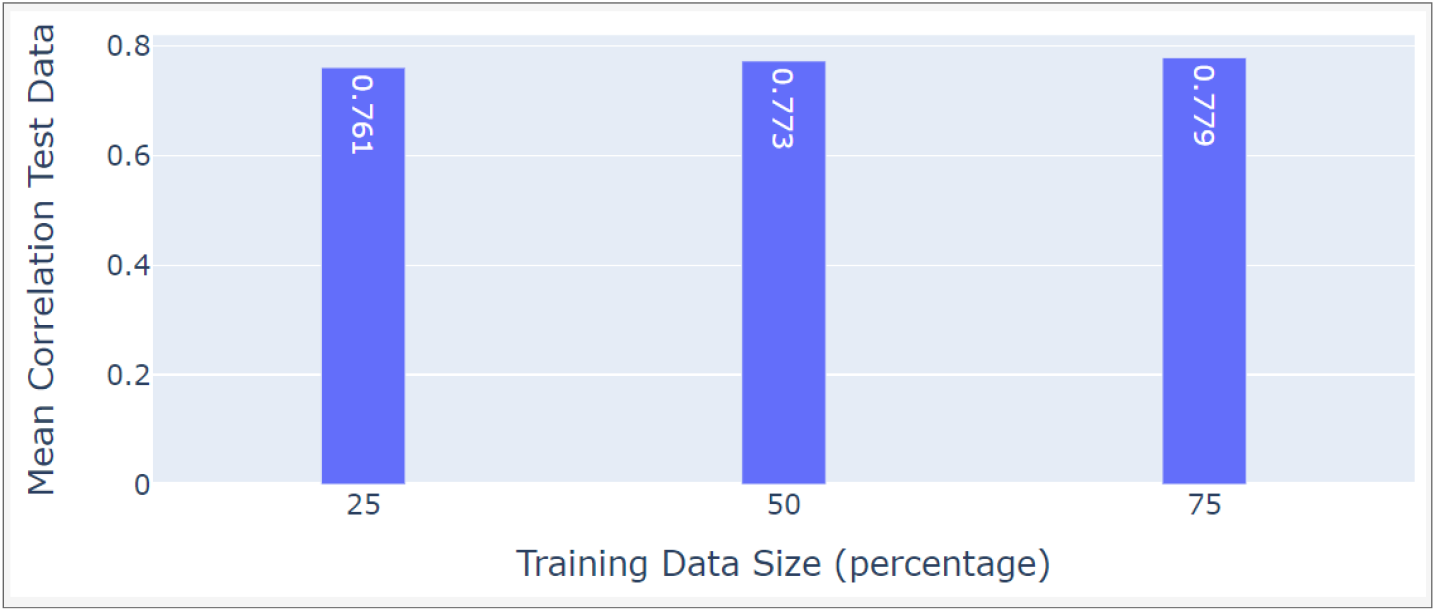
Effect of changing the training set size on the model performance.

#### Perturbation Experiments with Testing Dataset

We perturbed the data corresponding to the 48 test subjects from the 5-run experiment reported earlier, where each subject was perturbed *N* = 250 times. Here, each test SC matrix was perturbed by randomly generating the values of the elements from a powerlaw distribution that the elements are known to follow [38]. The A-GHN model was trained on unperturbed data of SC-FC pairs (50 subjects), and the resulting model was tested on each perturbed set of the test SC-FC pair. Figure 13 depicts the distribution of average Pearson correlation scores for these experiments. It can be observed that the model learned from the 50 unperturbed SCs performs rather poorly in predicting the FCs estimated from the randomly generated SCs. The histogram of mean correlation values ranges in [0.12, 0.45] with a mean correlation around 0.3, thus indicating that the model performance deteriorates significantly when fed with random structural connectivity information during the testing period. Thus, we can empirically conclude that the proposed model indeed learns SC-FC mapping, and the FC predictions are not independent of SC but respect the topology/structure of the input.

**Figure 13:**
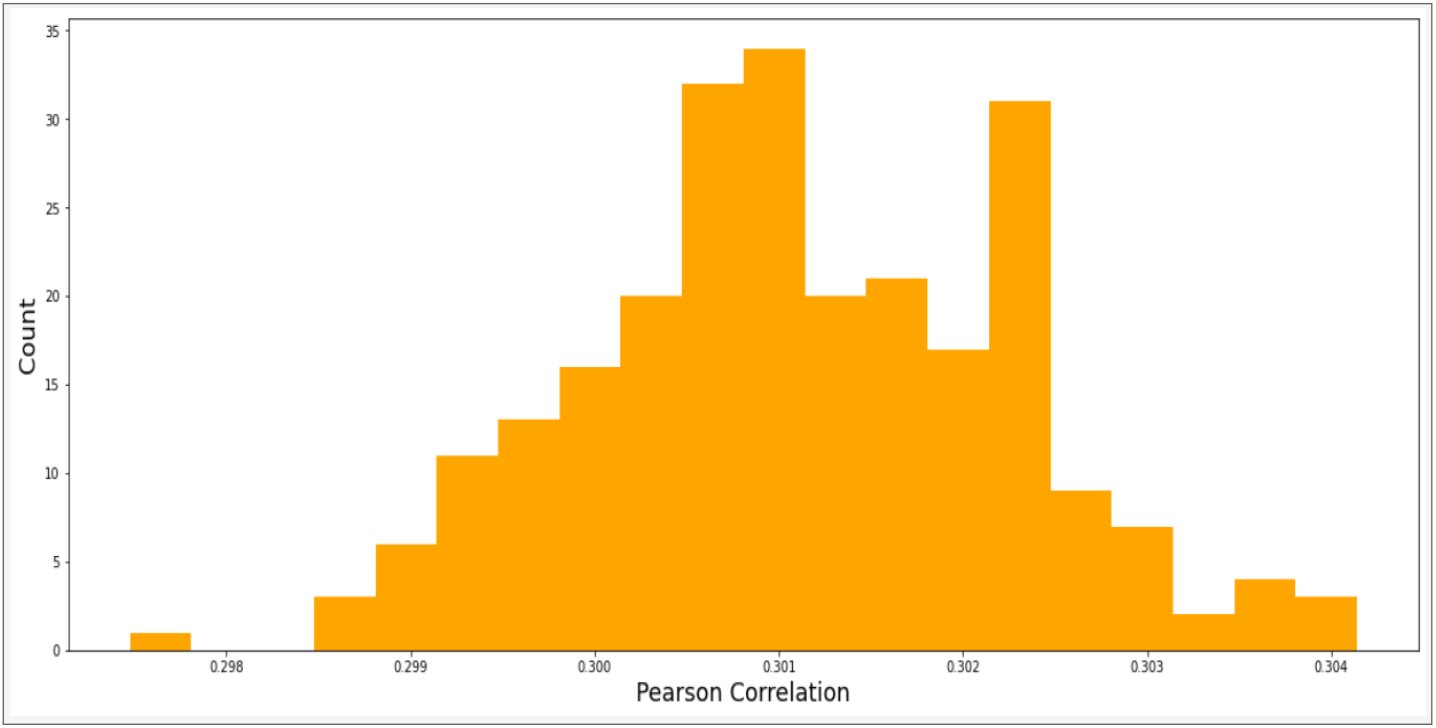
The Figure shows the average Pearson correlation scores distribution in the perturbation experiments related to the testing dataset. The correlation scores between the predicted FC estimated using randomly perturbed SCs (N = 250 sets) of the test subjects and the ground truth FC is considered here. Randomizing the SC inputs for the test dataset seems to impair the performance severely.

#### Perturbing the Model Input

In the second perturbation study, we trained the A-GHN model with perturbed SCs during the training phase and tested using the ground truth SC-FC pairs. Basically, this experiment tries to check the influence of training with broken training set on the target prediction ability. To assess whether FC prediction relies on the specific test SCs during the training phase, we considered the 50 random subjects used in the 5-run experiments reported earlier. Here, each training subject’s SC matrix was perturbed *N* = 250 times by randomly generating the values of the elements from a power-law distribution that the elements are known to follow [38]. The 250 perturbed sets of SCs for each training subject were used to train independent A-GHN models. These trained models were used for testing with the correct test SC-FC pairs (48 subjects). Figure S9 (in the Supplementary) depicts the histogram of the mean performances across all the sets. As observed, the mean correlation is around 0.34 with the correlation values ranging in [0.24, 0.41]. The poor performance indicates the importance of training with meaningful structurefunction relationships in order to yield valid predictions of the FC.

## 5. Discussion

The study of the relationship between structural connectivity and functional connectivity and how the functional activity of the brain is generated from the anatomical structure has been a major research topic in the field of cognitive neuroscience. Several methods have been proposed to explore the mapping between SC-FC including, whole brain computational models [22, 23], simple linear diffusion models [3] as well as complex non-linear models [25, 27], and linear multi-scale diffusion models [6, 7]. The whole brain computational models have been used as powerful tools to understand the relationship between structural and functional brain connectivity by linking brain function with its physiological underpinnings. On the other hand, non-linear complex drift-diffusion models based on excitatory and inhibitory neuronal populations, though not analytically tractable, give rise to rich dynamics. Abdelnour et al. [3] introduced a graph-based model with a linear single scale diffusion kernel at an optimal scale over the structural graph topology (SC) to map FC. However, Surampudi et al. [6] showed that single kernel models do not generalise to a larger cohort and demonstrated that FC can be decomposed into multiple diffusion kernels with subject non-specific combination coefficients. Further, the MKL framework, proposed by Surampudi et al. [7], revealed that the combination of multiple diffusion kernels was not sufficient to explain the self-organizing resting-state patterns found in FC and hence necessitated the use of additional explanatory parameters.

In this paper, we adopt the representation of the graph signal in terms of graph heat kernel similar to GraphHeat proposed by [16]. The GraphHeat formalism allows for selective focus on low-frequency spectral components at higher scales, whereas high-frequency spectral components are suppressed at lower scales. We consider a bank of such GHN models, each associated with a scale-specific heat kernel over the SC graph as input. The proposed A-GHN model then combines the outputs of the scale-specific GHN models using attention-based fusion. Both the hidden parameters (*W_γ_i__*) associated with the scale-specific GHN models as well as the attention scores that combine the A-GHN sub-model outputs are jointly learned to estimate the empirical FC accurately. We have established a correspondence between the initial regional co-activation parameters (*W_γ_i__*) in the proposed model and the parameters (Π_*i*_) from the MKL framework [7]. It is to be noted that the MKL framework is shown to be a variant of a reaction-diffusion system on the graph topology determined by the underlying structural connectivity (SC) matrix. Thus, the proposed A-GHN method is grounded in the theory of the reaction-diffusion process in the cognitive domain.

The proposed A-GHN model displays superior performance as compared to baseline models such as GCN Encoder-Decoder [15] and MKL model [7]. The model is able to learn population patterns regarding the SC-FC relationship even with smaller datasets. We validated our proposed model in three different settings: (i) 5-runs with the random initialization, (ii) LOO-CV, and (iii) 5-Fold cross-validation. The experimental results showed that the correlation structure of the BOLD functional resting-state brain networks is significantly well captured by our model (Fig. 4). The predicted mean correlation for 48 test subjects is close to 0.773 (5-Runs experiment), 0.803 (LOO-CV), whereas the GCN Encoder-Decoder and MKL yield (0.73, 0.74), and (0.69, 0.71), respectively. We conducted several ablation studies and perturbation experiments to establish the robustness of the reported results.

As explained below, the proposed framework enjoys three key properties of generalisability, scalability, and tractability in the deep learning framework.

### Interpretability & Generalisability

We formulate the deep learning model, A-GHN, as an end-to-end framework for SC-FC prediction. The challenge in applying deep learning models to neuroimaging research lies in the black-box nature of the process, where it is hard to decipher what the deep network actually learns. In order to address this and to understand the model mechanisms, we devised the following: (i) deciphering the learned parameters *W_γ_i__*, (ii) visualising the outputs of *m* number of A-GHN sub-models (Ψ_*γ_i_*_), and (iii) displaying the heatmap of attention probabilities across the test subjects (48 pairs of SC-FC), as shown in Figures 14, 15, and 16, respectively.

**Figure 14:**
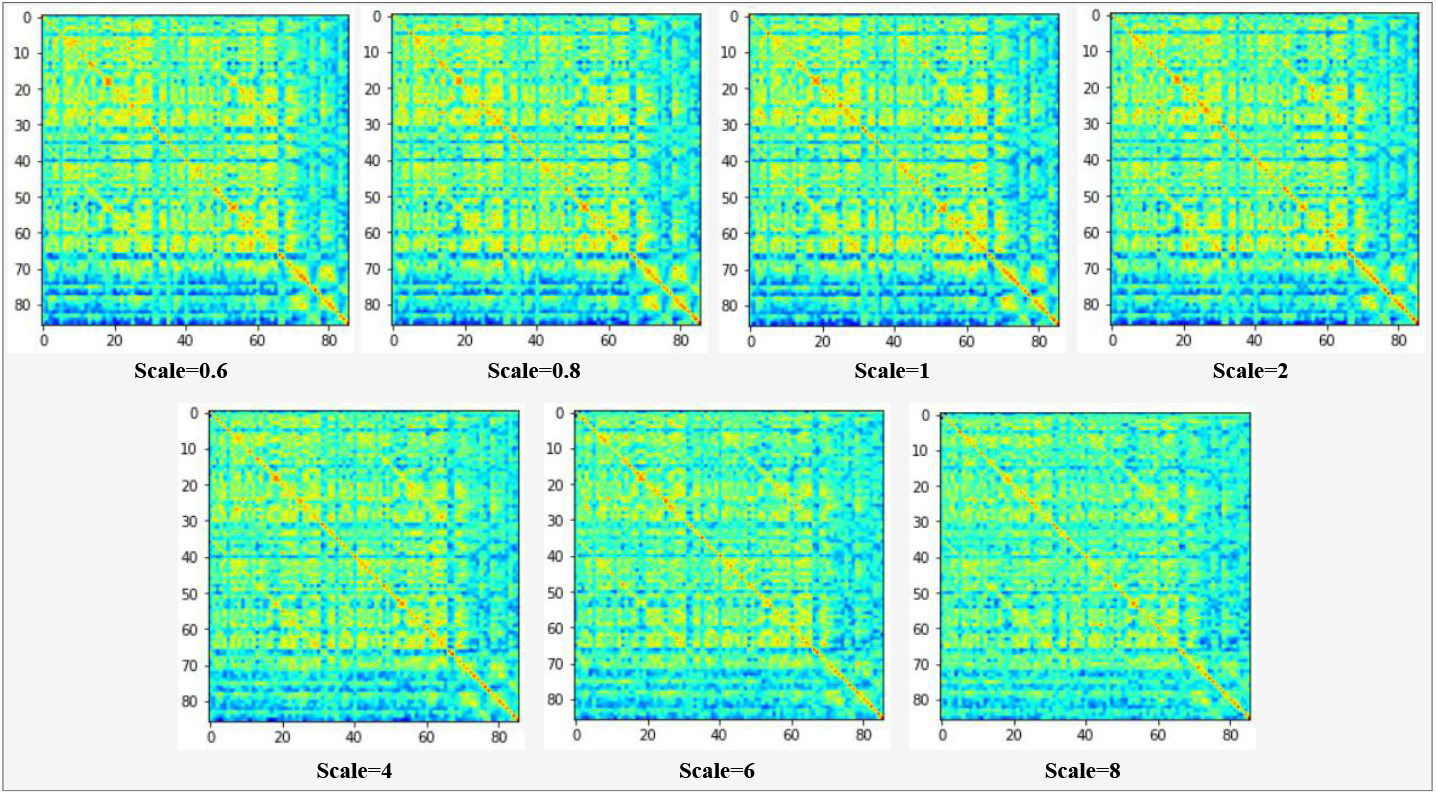
Distinctness of learned weights (*W_γ_i__*’s) corresponding to A-GHN sub-models. We check the distinctness of *W_γ_i__*’s for every scale value ranging from i = 1, ···, m (= 7). Each of these *m* matrices is of 86×86 dimension.

**Figure 15:**
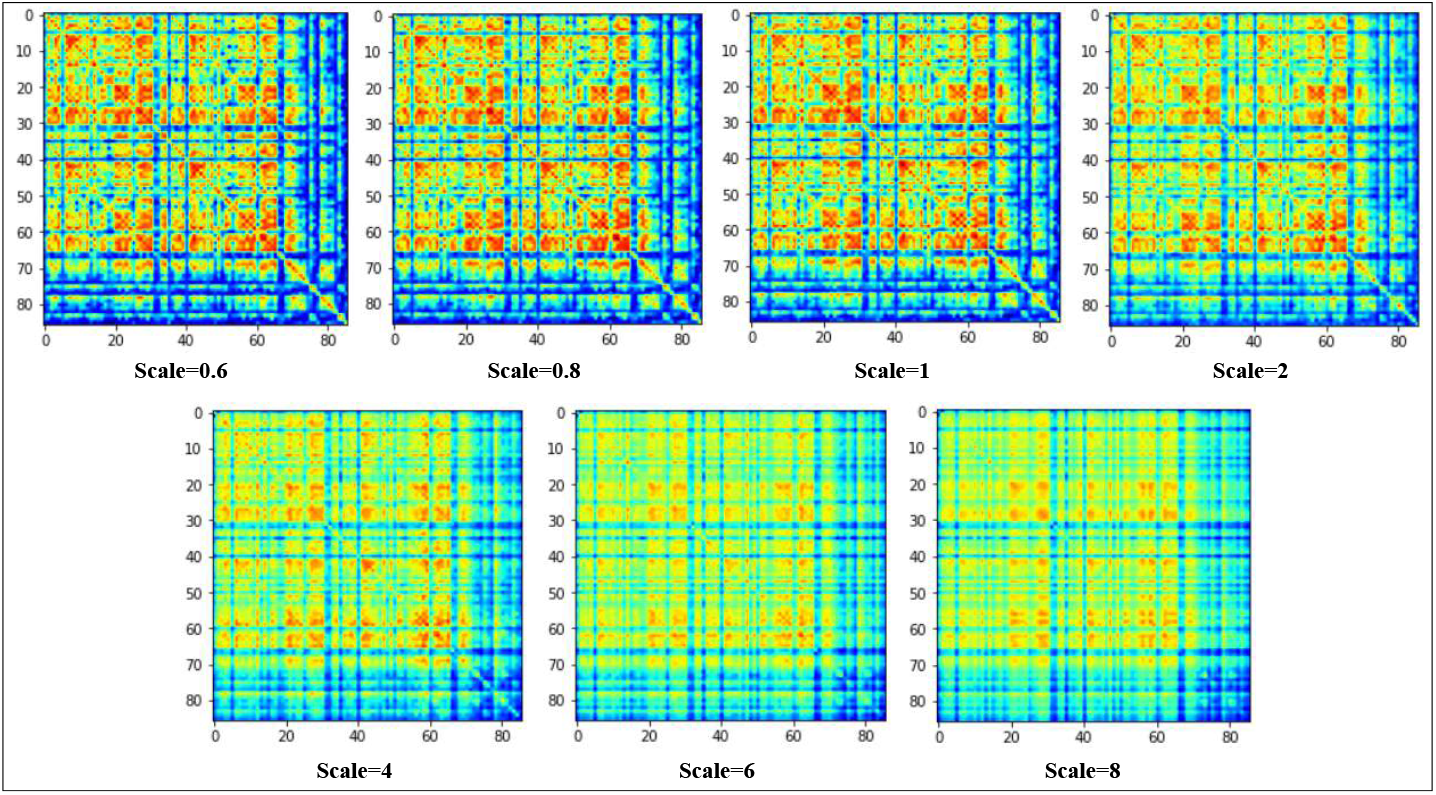
Distinctness of *ψ_γ_i__*’s. After scale-specific A-GHN sub-models are learned, we check the distinctness of *ψ_γ_i__*’s for every scale value ranging from i = 1,···, m (= 7). Each of these *m* matrices is of 86×86 dimension.

From Figures 14 & 15, we observe that lower scales display mean regional activity local to the neighboring nodes by suppressing the high-frequency spectral components. However, as the scale value increases, the large neighborhoods are taken into account with a global structure and captures much more information while discarding some irrelevant low-order neighbors. Thus, the proposed A-GHN model thereby be tuned to produce both local and global connectivity at lower and higher scales, respectively. Similarly, Figure 16 reports that the contribution of attention probabilities is decreasing as the scale value increases. Further, we performed community detection to identify the different networks captured in the FC predicted by the model. The communities were detected using the Louvain algorithm as described in the Brain Connectivity Toolbox(BCT) [36]. From Figure 17 it is observed that the communities detected in the predicted FC when compared with empirical FC(ground truth) captures inter-hemispherical patterns as well. We also showcase the communities detected in each scale-specific sub-models of A-GHN, as shown in Figure S6 (Refer Supplementary). The current work demonstrates the feasibility of the A-GHN model with experiments on a medium-size dataset of 100 participants. However, the generalisability of larger datasets can be explored in the future.

**Figure 16:**
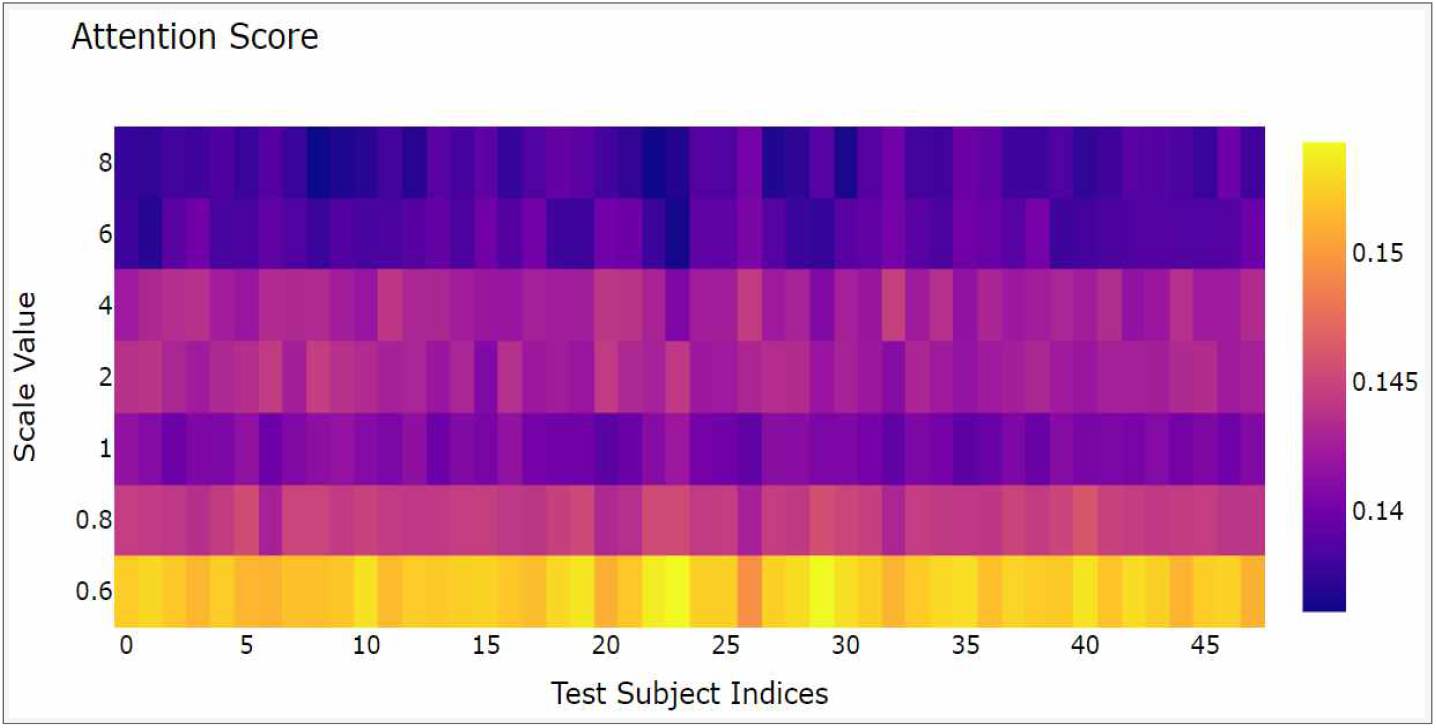
The Figure showcase the contribution of the attention scores associated with each scale-specific A-GHN sub-model output related to a testing dataset in the A-GHN model. The attention score probabilities for the smaller scales are marginally higher than large-scale.

**Figure 17:**
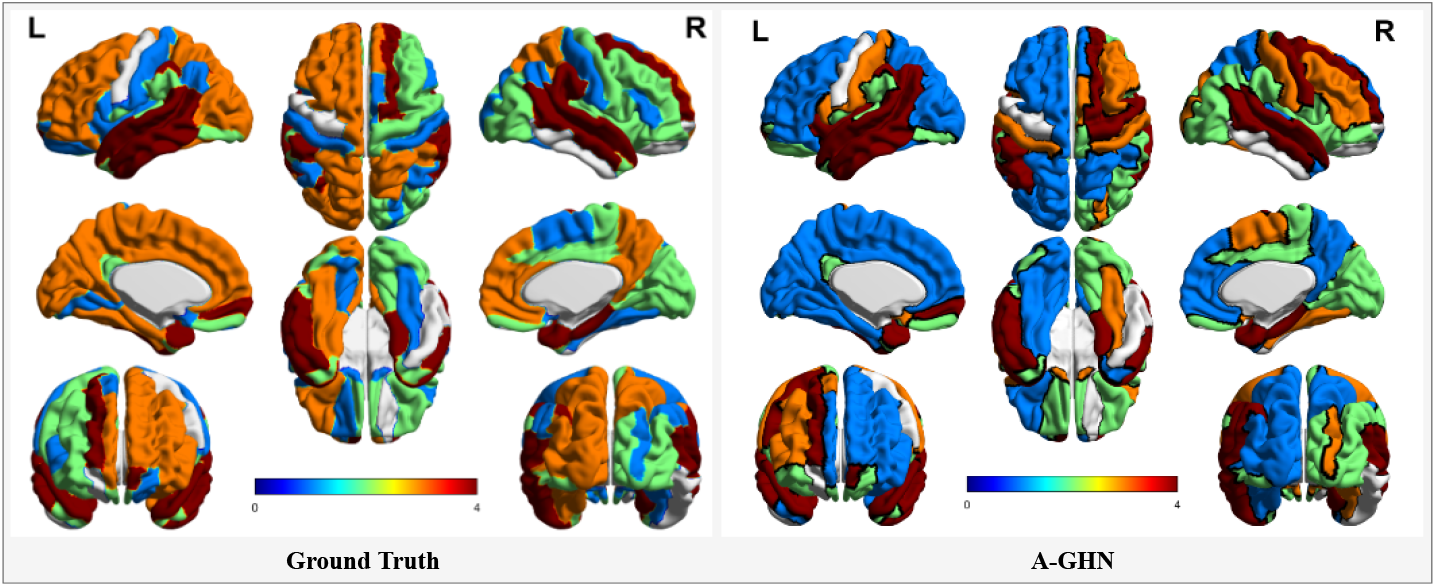
The distinct, modular communities uncovered in empirical FCs (Ground Truth) and predicted FCs using the A-GHN model. The communities detected using BCT [36] are mapped onto the brain surface using BrainNet Viewer [37]. The colors are mapped according to the communities uncovered by the stochastic Louvain algorithm defined in BCT. Thus, the color scale cannot be directly compared between ground truth and A-GHN.

### Scalability & Computational Efficiency

The results reported in the current work use the parcellation based on the AAL Atlas (86 × 86). Nevertheless, the A-GHN model is easily scalable to any brain parcellation (for example, Gordon Atlas with 333 × 333, Glasser Atlas with 360 × 360 parcellations). Graph-based diffusion models [3, 9, 7, 39] that are not easily scalable for larger parcellations as the matrix operations are difficult to scale for larger matrix sizes. On the other hand, since graph convolutional network (GCN)-based models [15] including the proposed A-GHN model use only node aggregate features that require vector operations; hence they are easily scalable.

From a computational efficiency perspective, one of the major limitations of the MKL model [7] is that it uses LASSO optimization that requires computationally expensive matrix inverse operations. Hence the computational complexity is dominated by the cost of LASSO optimization. In contrast, the proposed A-GHN model is more efficient as it uses a stochastic gradient-based backpropagation learning approach. Moreover, the A-GHN model requires learning of 59,168 parameters (7 scales: 7×7396 + Attention Module: 1×7396) that is comparatively lower than learning 118,336 parameters in the MKL framework (16 scales: 16×7396). Further, the proposed framework is inherently scalable to more diffusion scales, more hidden layers in the GHNs, and can potentially be used for transfer learning on other datasets – all these make the A-GHN model very flexible and computationally powerful.

### Limitations & Future Work

Usually, deep learning models require large datasets to obtain reliable learning and generalisation performance results. An interesting point to note of our work is that it is trained and tested on a medium-size dataset. We demonstrated how A-GHN can be trained to obtain superior results using hyperparameter tuning and various validation experiments even with such a dataset. It would be interesting to demonstrate how A-GHN scales to larger datasets in the future. This research is the first step in applying the A-GHN model to perform automatic resting-state FC prediction from SC. In the near future, we intend to use the A-GHN model as a universal model to predict the FC of different types (both resting-state FC as well as task-based FCs) with the structural graph given as input.

In future work, a biophysical interpretation of the proposed deep learning model (A-GHN) with multi-scale heat kernel diffusion as an instance of a reaction-diffusion system on the structural brain graph needs to be established. Additionally, the proposed model could be used to characterize the disease groups as well. A-GHN considers average functional connectivity, ignoring the transient functional dynamics over the period of acquisition of the temporally extended rsfMRI signal. The proposed framework could potentially be extended to capture the temporal information in the functional connectivity dynamics (FCD).

## 6. Conclusion

This paper proposed a novel A-GHN model that outperforms existing models that use either multiple diffusion kernels (MKL) or that use GCNs (GCN Encoder-Decoder). Extensive cross-validation, perturbation, and ablation studies establish the robustness of the proposed architecture for learning the structure-to-function mapping of the brain using the images from DTI and rsfMRI. The model not only captures the SC-FC mapping but the underlying functional connectivity networks as well. The strengths of the deep learning based GHN models over graph diffusion-based linear models such as the MKL model are their computational efficiency and scalability.

